# The Genetic Equidistance Phenomenon at the Proteomic Level

**DOI:** 10.1101/031914

**Authors:** Denghui Luo, Shi Huang

## Abstract

The field of molecular evolution started with the alignment of a few protein sequences in the early 1960s. Among the first results found, the genetic equidistance result has turned out to be the most unexpected. It directly inspired the ad hoc universal molecular clock hypothesis that in turn inspired the neutral theory. Unfortunately, however, what is only a maximum distance phenomenon was mistakenly transformed into a mutation rate phenomenon and became known as such. Previous work studied a small set of selected proteins. We have performed proteome wide studies of 7 different sets of proteomes involving a total of 15 species. All 7 sets showed that within each set of 3 species the least complex species is approximately equidistant in average proteome wide identity to the two more complex ones. Thus, the genetic equidistance result is a universal phenomenon of maximum distance. There is a reality of constant albeit stepwise or discontinuous increase in complexity during evolution, the rate of which is what the original molecular clock hypothesis is really about. These results provide additional lines of evidence for the recently proposed maximum genetic diversity (MGD) hypothesis.

**Availability and implementation:** The source code repository is publicly available at https://github.com/Sephiroth1st/EquidistanceScript

**Supplementary information:** Supplementary data are available online.

## 1 Introduction

The field of molecular evolution first started in the early 1960s (Doolittle and Blombaeck, 1964; Margoliash, 1963; Zuckerkandl and Pauling, 1962). For any three or more species of different organismal complexity as intuitively defined by the number of cell types, one can perform two kinds of sequence alignment. The first aligns a complex organism such as human against simpler or less complex species that evolved earlier such as frogs and fishes. The second aligns simpler organisms such as fishes against those more complex ones such as frogs and humans.

The first kind of alignment was first done using hemoglobins and showed the pattern that human shares more identity with mammals than with fishes, which is largely consistent with Darwinian expectations (Zuckerkandl and Pauling, 1962). Margoliash in 1963 performed both alignments and made a formal statement of the molecular clock after noticing the genetic equidistance result: “It appears that the number of residue differences between cytochrome c of any two species is mostly conditioned by the time elapsed since the lines of evolution leading to these two species originally diverged. If this is correct, the cytochrome c of all mammals should be equally different from the cytochrome c of all birds. Since fish diverges from the main stem of vertebrate evolution earlier than either birds or mammals, the cytochrome c of both mammals and birds should be equally different from the cytochrome c of fish. Similarly, all vertebrate cytochrome c should be equally different from the yeast protein.” (Kumar, 2005; Margoliash, 1963).

In hindsight, however, Margoliash could as well have proposed an alternative interpretation: “It appears that the number of residue differences between cytochrome c of any two species is mostly conditioned by the species with lower organismal complexity. If this is correct, the cytochrome c of all mammals should be equally different from the cytochrome c of all birds. Since fish has lower complexity than either birds or mammals, the cytochrome c of both mammals and birds should be equally different from the cytochrome c of fish. Similarly, all vertebrate cytochrome c should be equally different from the yeast protein.” The fact that he did not was unfortunate for the field as it mistakenly converted a maximum distance phenomenon into a rate phenomenon.

The constant mutation rate interpretation of the equidistance result has in fact turned out to be a classic tautology since it has not been verified by any independent observation and has on the contrary been contradicted by a large number of facts (Avise, 1994; Goodman, et al., 1974; Huang, 2008; Huang, 2008; Huang, 2009; Huang, 2009; Jukes and Holmquist, 1972; Laird, et al., 1969; Langley and Fitch, 1974; Li, 1997; Nei and Kumar, 2000; Pulquerio and Nichols, 2007). Nonetheless, researchers have treated the molecular clock as a genuine reality and have in turn proposed a number of theories to explain it (Clarke, 1970; Kimura, 1968; Kimura and Ohta, 1971; King and Jukes, 1969; Richmond, 1970; Van Valen, 1974). The ‘Neutral Theory’ has become the favorite (Kimura, 1968; Kimura and Ohta, 1971; King and Jukes, 1969), even though it is widely acknowledged to be an incomplete explanation for the clock (Ayala, 1999; Pulquerio and Nichols, 2007). The observed rate is measured in years but the Neutral theory predicts a constant rate per generation. Also, the theory predicts that the clock will be a Poisson process, with equal mean and variance of mutation rate. Experimental data have shown that the variance is typically larger than the mean.

Ohta’s “nearly neutral theory” explained to some extent the generation time issue by observing that large populations have faster generation times and faster mutation rates but remains unable to account for the great variance issue (Ohta, 1973). With the neutral and nearly neutral theory, molecular evolution has been treated as the same as population genetics. However, the field still lacks a complete theory as Ohta and Gillespie had acknowledged: “We have yet to find a mechanistic theory of molecular evolution that can readily account for all of the phenomenology. Thus, while the 1990s will most likely be a decade dominated by the gathering of data, we would like to call attention to a looming crisis as theoretical investigations lag behind the phenomenology.” (Ohta and Gillespie, 1996). The field has unfortunately yet to pay attention to the equidistance result, which has been considered by some as “one of the most astonishing findings of modern science”(Denton, 1986).

We recently proposed the maximum genetic diversity hypothesis based on a pair of intuitions or axioms (Hu, et al., 2013; Huang, 2008; Huang, 2009; Huang, 2016). Axiom 1 says that the more complex the phenotype, the greater the restriction on the choice of molecular building blocks. Axiom 2 says that any system can allow a limited level of random errors or noises in molecular building parts and such errors may be beneficial, deleterious, or neutral depending on circumstances. Obviously, one only needs to substitute “errors in molecular building blocks” for “genetic diversity” to get the equivalent concept in biology. Axiom 2 in effect, in our opinion, highlights the proven virtues of the modern evolution theory consisting of Darwin’s and Kimura’s theories.

Genetic diversity or distance cannot increase indefinitely with time and has a maximum limit being restricted by function or epigenetic complexity. The maximum genetic diversity of simple organisms is greater than that of complex organisms. Over long evolutionary time, the genetic distance between sister species and a simpler outgroup taxon is mainly determined by the maximum genetic diversity of the simpler outgroup, although over short time scales it is mainly determined by time, drift, environmental selection, and the neutral mutation rates of the simpler outgroup as well as to a smaller extent by the rates of the sister taxa. The MGD hypothesis thus includes the proven virtues of modern evolution theory, consisting of Darwin’s theory and the neutral theory, as relevant only to microevolution over short time scales before sequence divergence reaches maximum genetic distance/diversity. In other words an increase in epigenetic complexity during macroevolution is associated with a suppression of genetic diversity or point mutations.

The MGD hypothesis explains the genetic equidistance phenomenon as a result of maximum genetic distance imposed by epigenetic constraints (Hu, et al., 2013; Huang, 2008; Huang, 2009). This phenomenon has in fact another characteristic, the overlap feature where particular sites in an amino acid sequence are subject to multiple different mutational changes in a particular lineage, which has been overlooked for nearly half of a century (Huang, 2010). Overlapped mutant amino acid positions are detected where any pair of any three species is different, which indicates repeated turnover of residues at the same position. These overlap positions exhibit a strikingly non-random pattern in complex organisms indicative of severe epigenetic constraints during macroevolution. In contrast, when simple organisms of similar complexity and of short evolutionary divergence are compared, there are only a small number of overlaps largely consistent with chance or the neutral theory, indicating a random distribution of mutations during microevolution. So, while the molecular clock may superficially explain the apparent equi-distance in numbers, it cannot explain the non-random distribution of mutation hot spots and the related observation that the percentage of constrained sites in more complex clades is greater than that in simpler organisms (Huang, 2010).

The MGD hypothesis has accounted for most major phenomenology of molecular evolution. It has also been instrumental in directing productive research into not only evolutionary phylogenetic problems but also important biomedical problems of today, including the genetic understanding of complex traits and diseases (Huang, 2012; Huang, 2016; Yuan, et al., 2012; Yuan, et al., 2014; Zhu, et al., 2015; Zhu, et al., 2015; Zhu, et al., 2015; Zhu, et al., 2015). That human is closer to chimpanzees than to orangutans in genome wide identity is in fact due to functional or physiological similarity (hence the related molecular similarity) rather than more recent common ancestry (Huang, 2012). Orangutan has less reasoning ability than chimpanzee and human (Herrmann, et al., 2007). Therefore, the rationale for genetic equidistance equally explains the increased amount of difference found when human proteins are compared with homologs from progressively more distant or progressively less complex species.

The equidistance result, as long as it only means approximate, appears to be a feature of almost all proteins as indicated by a random sampling of 50 proteins (Huang, 2008). Others have presented this in passing in 20 most conserved proteins (Copley, et al., 1999). Recent advances in whole genome sequencing have made it possible to examine whether this phenomenon holds at the proteome level.

## 2 Methods

### 2.1 Protein datasets

Protein sequence data were downloaded from UniProt (http://www.uniprot.org), which has collected 2404 proteomes from various organisms.

### 2.2 Software

Analyses were performed on MacOS using BASH. Sequence alignments used downloaded BLAST 2.2.29+(Shiryev, et al., 2007), and Clustal Omega -1.2.0(AndreaGiacomo) (Sievers, et al., 2011).

We wrote a few custom scripts as follows:

> blastp3.sh for protein sequence alignment of three species (running this script need blast+ program and getHomGap.pl script).
>
> calWeightPident.awk to calculate weighted identity taking into consideration of protein length.
>
> getHomGap.pl to identify orthologs based on reciprocal matches of a pair of species.
>
> getEach3OrthologProtein.pl to extract each alignment of 3 orthologs set.
>
> overlapsCal.pl to calculate the overlap feature in a three species alignment.

### 2.3 Analysis of genetic equidistance at proteomic level

To study alignment of an individual protein conserved in multiple species, we obtained its sequence from multiple species and used clustal omega to obtain a multi-species alignment. The percentage identity between each pair of species was scored using BLASTP.

To study genetic equidistance of proteomes, we picked groups of three species each group for our study. We picked out orthologs conserved in all three species by doing pairwise alignments. Whole proteome data in fasta format were downloaded and built as a library suitable for BLASTP analysis by using mkblastdb of the BLAST+ software. Pairwise whole proteome alignments were then performed using BLAST. The results were cleaned by removing redundant alignments to retain only those between candidate orthologs. We identified candidate orthologs based on bitscore results of BLAST by following the reciprocal best hit method (Moreno-Hagelsieb and Latimer, 2008). The basic procedure entails collecting all the genes in two species and comparing them to one another. If genes from two species identify each other as their closest partners then they are considered orthologs. This works well for closely related species but can be a major problem in highly divergent species. There is a tradeoff between specificity and coverage. For our selection, we restricted the length of the orthologs to be a minimum of 100 amino acids. We aimed for specificity instead of coverage.

We then merged the alignment results of pairwise comparisons into an alignment of three species. We wrote a BAST script blastp3.sh for performing the whole process from the fasta sequence data to the final three species alignments. The script wil invoke other related scripts makeblastdb, blastp and getHomGap.pl.

### 2.4 Scoring overlapped positions

We picked three species, zebrafish, xenopus, and human, to study the overlap feature. We selected the top 50 ranked proteins in BLAST alignment bitscore. From these, we picked 7 most conserved and 7 least conserved and put the names of each protein in a new document. We then used the custom script getEach3OrthologProtein.pl to pick out the protein sequence from the library to form a new fasta document, which was next used by the overlapsCal.awk script to calculate the overlap ratio. The overlap ratio is defined as the number of actual overlap positions divided by the number of candidate positions (Huang, 2010). The candidate overlap positions in any three species comparison involving an outgroup include all the different positions between the two sister lineages.

### 2.5 Statistics methods

The degree of identity between two orthologs proteins was weighted by sequence length. The longer the protein the higher the weight score. We wrote the CalWeightPident.awk script to do this. A weighted identity score for each protein was calculated by the formula: (percent identity x protein length) / average length of all proteins. The weighted identity score with standard deviation was used for F test and Student’s t test. Pearson and Spearman analyses were performed to examine the correlation between overlap ratio and protein conservation.

## 3 Results

### 3.1 Genetic equidistance in multiple species

Genetic equidistance in multiple (>3) species alignment has only been shown for cytochrome C, one of the three proteins first analyzed (Doolittle and Blombaeck, 1964; Margoliash, 1963; Zuckerkandl and Pauling, 1962). We here generated another example with the Dot1 His-tone-lysine N-methyltransferase (H3 lysine-79 specific). The selection of this protein was for no particular reason other than our past research interest in the area of protein methylation and the highly conserved nature of this protein which makes multiple species alignments possible.

We aligned the shared portions of Dot1 orthologs from 8 species (Supplementary Figure S1) and obtained the percentage difference values (Table 1). As is evident when each column in Table 1 is viewed vertically, each apparently low complexity species (as judged by common sense and the estimated number of cell types) is approximately equidistant to all higher complexity species. For example, yeast is ∼70% different from worm, sea urchin, zebrafish, frog, zebra finch, bat, and human. Fish is ∼8% different from frog, zebra finch, bat, and Human. When the rows in Table 1 are viewed horizontally, a gradual decrease from left to right in protein non-identity is obvious and accompanies the increase in organismal complexity.

**Table 1.**
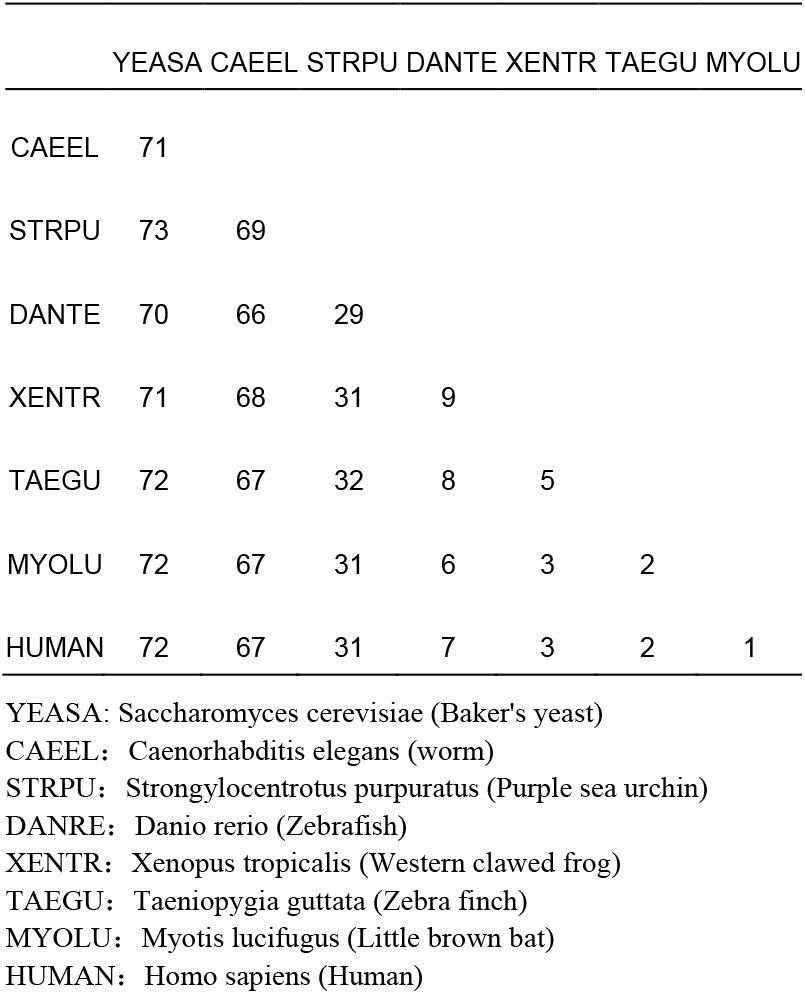
Percentage difference among species in Dot1.

### 3.2 Genetic equidistance in multiple sets of three proteomes comparisons

To analyze genetic equidistance at the proteome level, a minimum of three species is needed where two sister species could be compared to an outgroup to determine whether they are equidistance to the outgroup. Here the concept of sister species is not about one’s closest sister but is relative to the outgroup. In order to examine the universal nature of the equidistance phenomenon, we would like to select appropriate species that could cover a wide range of the biological world. We selected the following: a bacteria Escherichia coli (strain K12), a fungi Saccharomyces cerevisiae (strain AWRI796),a protozoa Paramecium tetraurelia, a worm Caenorhabditis elegans, an insect Apis mellifera (The western honey bee), a sea urchin Strongylocentrotus purpuratus, a fish Danio rerio (Zebrafish), an amphibian Xenopus tropicalis (Western clawed frog), a lizard Anolis carolinensis, a duck Anas platyrhynchos), a bird Taeniopygia guttata (Zebra finch), a primitive mammal Duckbill platypus Ornithorhynchus anatinus, a flying mammal Myotis lucifugus (Little brown bat), a monkey Callithrix jacchus (White-tufted-ear marmoset), and four hominoids Gorilla gorilla, Pongo abelii (Sumatran orangutan, Pan troglodytes (Chimpanzee), and homo sapiens.

From these species, we designed 7 sets of three species combinations where the three species are of different apparent complexity. We expect to see the least complex outgroup to be equidistant to the other two sister species. 1. To test equidistance to bacteria, we picked bacteria, yeast, and human. Most would naïvely expect bacteria to be closer to yeast than to human. 2. To test equidistance to protozoa, we selected protozoa, worm, and insect. Protozoa is unicellular eukaryotes and expected to be equidistant to multicellular organisms. Worm seems less complex than insect. 3. To test equidistance to fish, we selected fish, amphibian, and human. Fish is the least complex and expected to be equidistant to frog and human. 4. To test equidistance to amphibian, we selected amphibian, lizard, and human. 5. To test equidistance to a bird, we selected duck, platypus, and human. Duck seems the least complex among the three. 6. To test equidistance to monkey, we tested monkey, gorilla and human with monkey the least complex. 7. To test equidistance to a hominoid, we selected orangutan, chimpanzee, and human. Orangutan has less reasoning ability than chimpanzee and human (Herrmann, et al., 2007).

We selected orthologs of minimum of 100 amino acids in length for sequence comparison. For each of the seven sets, the average alignment gaps between each pair of orthologs within the set were similar (Supplementary Table S1-S7). We obtained the percentage identity for each protein. We also calculated a weighted identity score for each protein by the formula: (percent identity x protein length) / average length of all proteins. The weighted score is more suitable for a meaningful average protein identity as a longer length protein should contribute more to the average than a short one.

The average pairwise identity and weighted identity score among the orthologs in each set are shown in Table 2 and Supplementary Table S1-S7. The results were all consistent with expectations of equidistance to the less complex outgroup species. For example, proteome average identity between bacteria and yeast is 34.3%, similar to 35.8% between bacteria and human. After adjustment with more weight on longer length proteins, the result became more obvious with identity between bacteria and yeast at 35.2%, and identity between bacteria and human at 36% (P = 0.76, t test, Table 4). Similar results were found for all 7 sets of comparisons (Table 2 and 3).

**Table 2.** Average pairwise identity, protein length, number of gaps, and weighted identity of orthologs from different species.

| Species |  |  | Average Identity |  |  | Average Length |  |  | Average number of Gaps |  |  | Weighted Average Identity |  |  | Protein Number |
| --- | --- | --- | --- | --- | --- | --- | --- | --- | --- | --- | --- | --- | --- | --- | --- |
| 1 | 2 | 3 | 1-2 | 1-3 | 2-3 | 1-2 | 1-3 | 2-3 | 1-2 | 1-3 | 2-3 | 1-2 | 1-3 | 2-3 |  |
| ECOL | YEAS | HOMO | 37.02 | 38.69 | 43.90 | 325.96 | 335.87 | 361.93 | 7.04 | 6.40 | 6.88 | 38.08 | 39.27 | 44.07 | 196 |
| PARA | ELEG | BEE | 39.30 | 40.74 | 50.01 | 380.72 | 388.23 | 441.78 | 7.67 | 7.46 | 6.40 | 38.38 | 39.64 | 48.11 | 597 |
| FISH | FROG | HOMO | 64.58 | 64.96 | 69.71 | 553.81 | 567.10 | 576.14 | 6.58 | 6.34 | 5.19 | 63.40 | 64.13 | 68.84 | 8450 |
| FROG | ANOL | HOMO | 69.37 | 68.74 | 73.19 | 562.74 | 589.06 | 591.76 | 5.47 | 5.52 | 4.77 | 68.58 | 67.99 | 72.62 | 9228 |
| DUCK | PLAT | HOMO | 75.03 | 75.18 | 77.51 | 500.58 | 591.28 | 533.76 | 3.78 | 4.17 | 3.44 | 74.82 | 74.67 | 76.91 | 7739 |
| MARM | GORI | HOMO | 92.32 | 93.31 | 97.76 | 554.56 | 569.09 | 570.75 | 1.28 | 1.00 | 0.53 | 92.57 | 93.62 | 97.70 | 13706 |
| ORAN | CHIM | HOMO | 96.59 | 97.06 | 98.46 | 546.15 | 558.33 | 575.42 | 0.77 | 0.61 | 0.36 | 96.67 | 97.11 | 98.51 | 14408 |

ECOL: *Escherichia coli* (strain K12) YEAS: *Saccharomyces cerevisiae* (strain AWRI796) PARA: *Paramecium tetraurelia* (protozoa) ELEG: *Caenorhabditis elegans* (worm) BEE: *Apis mellifera* (The western honey bee) FISH: *Danio rerio* (Zebrafish) FROG: *Xenopus tropicalis* (Western clawed frog) ANOL: *Anolis carolinensis* (Green anole) DUCK: *Anas platyrhynchos* (Mallard) PLAT: *Ornithorhynchus anatinus* (Duckbill platypus) MARM: *Callithrix jacchus* (White-tufted-ear marmoset)
GORI: *Gorilla gorilla gorilla* (Western lowland gorilla) ORAN: *Pongo abelii* (Sumatran orangutan) CHIM: *Pan troglodytes* (Chimpanzee) HOMO: *Homo sapiens*

**Table 3.** Results of Student’s t test of weighted identities.

|  | P value |
| --- | --- |
| ECOL-YEAS vs ECOL-HOMO | 0.76 |
| ECOL-YEAS vs YEAS-HOMO | 0.034 |
| YEAS-HOMO vs ECOL-HOMO | 0.056 |
| PARA-ELEG vs PARA-BEE | 0.75 |
| PARA-ELEG vs ELEG-BEE | 0.00027 |
| ELEG-BEE vs PARA-BEE | 0.0008 |
| FISH-FROG vs FISH-HOMO | 0.47 |
| FISH-FROG vs FROG-HOMO | 2.16E-05 |
| FROG-HOMO vs FISH-HOMO | 0.00047 |
| FROG-ANOL vs FROG-HOMO | 0.79 |
| FROG-ANOL vs ANOL-HOMO | 8.55E-05 |
| ANOL-HOMO vs FROG-HOMO | 2.68E-05 |
| DUCK-PLAY vs DUCK-HOMO | 0.68 |
| DUCK-PLAY vs PLAY-HOMO | 0.17 |
| PLAY-HOMO vs DUCK-HOMO | 0.39 |
| MARM-GORI vs MARM-HOMO | 0.29 |
| MARM-GORI vs GORI-HOMO | 9.29E-07 |
| GORI-HOMO vs MARM-HOMO | 0.00012 |
| ORAN-CHIM vs ORAN-HOMO | 0.86 |
| ORAN-CHIM vs CHIM-HOMO | 0.045 |
| CHIM-HOMO vs ORAN-HOMO | 0.069 |

**Table 4.**
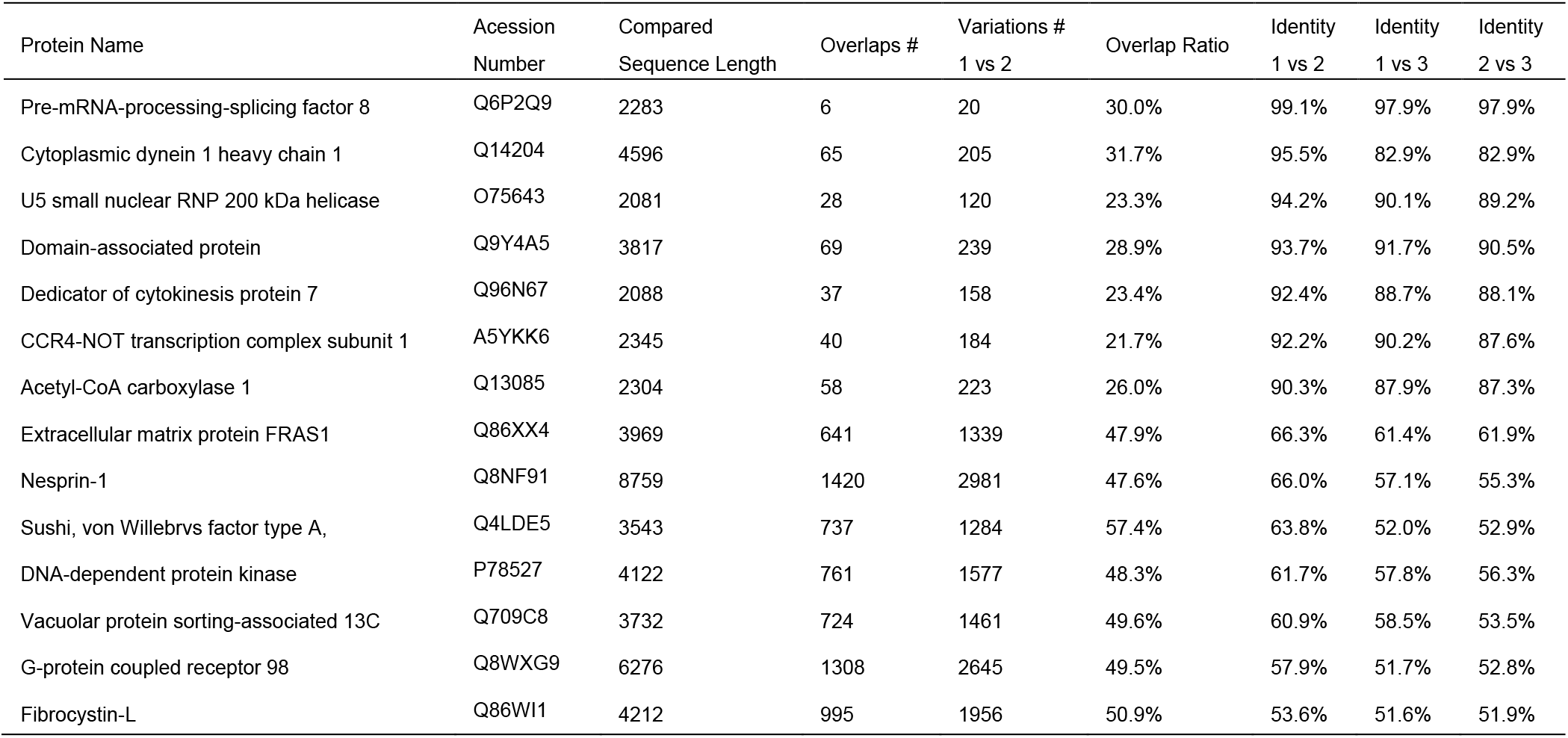
Overlap feature of selected proteins from human, frog, and fish with species represented by 1, 2, and 3 respectively.

### 3.3 The overlap ratio and protein conservation

The overlap feature has been studied for several proteins such as cytochrome C and hemoglobin (Huang, 2010). To further examine the universality of this feature, we here studied 7 fast evolving or low identity and 7 slow evolving or high identity proteins from the set of fish, frog and human. They were selected based on their quality of alignment (low number of gaps per unit length). As the proteins are selected from the top 50 ranked proteins in BLAST alignment bitscore, they should be relatively long in length. We counted the number of overlapped and candidate position and calculated the overlap ratio as the number of overlapped positions divided by the number of candidate positions (Table 4 and Fig. 1). The results showed significantly different overlap ratios for the set of slow evolving proteins versus the set of fast evolving proteins. The overlap ratio is inversely related to pairwise percentage identity between the two sister groups (Fig. 1).

**Figure 1.**
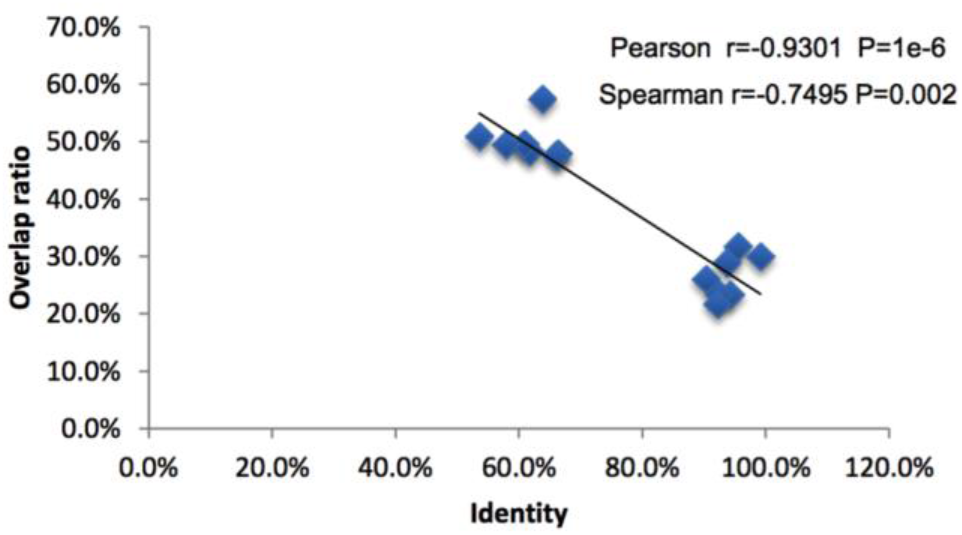
Relationship between the overlap ratio and the identity of the two sister species. Data in Table 4 were plotted and analyzed by Pearson and Spearman correlation tests.

## 4 Discussion

The genetic equidistance results have been previously found in a limited number of proteins and received little attention ever since the result was interpreted by the universal molecular clock hypothesis. Today, the universal nature of the molecular clock has been discredited by numerous findings of different mutation rates among different species. However none of those findings are based on the genetic equidistance phenomenon. Most researchers automatically assumed no equidistance whenever they found a non-constant rate. Few realized that the equidistance is not a result of constant mutation rates at least for fast evolving proteins that have reached MGD. Our results here show that the equidistance phenomenon is not merely specific to certain limited number of proteins but actually holds for nearly the complete proteome as a whole.

As our analysis involved a large number of species selected for no better reason than the availability of proteome sequences, the results indicate the universality of genetic equidistance with regard to species. The universal nature of this phenomenon in terms of both proteomes and species explains why the first result in protein alignment or molecular evolution is in fact the genetic equidistance result. One simply cannot miss it so long one is doing protein alignments.

For nearly half of a century, there have been only two scientific hypotheses to explain the genetic equidistance phenomenon, the molecular clock and the MGD hypothesis. Both are correct in their respective domains. As explained in previous publications, there are two kinds of equidistance, linear and maximum (Hu, et al., 2013). Over long evolutionary time scales or for fast-evolving sequences, maximum genetic equidistance is the predicted outcome: different species are equidistant to a species of lower or equal complexity, and such distances do not change with time. The sequential divergences between the cytochromes initially reported by Margoliash are a measure of maximum genetic equidistance (Margoliash, 1963). And the vast majority of examples of equidistance observed today are maximum since most sequences are non-coding or fast evolving, and have undergone evolution over long periods of time. For short evolutionary time scales or for slow-evolving sequences, linear genetic equidistance is the norm where the molecular clock holds and the distance is still linearly related to time: when sister species have similar mutation rates, they would be equidistant to a less or equally complex outgroup, and such distances still increase with time.

The two kinds of equidistance can be easily distinguished by the overlap feature (Huang, 2010). Where the equidistance has reached a maximum a large overlap ratio is observed while linear equidistance has none or few overlap positions. Unfortunately, the field has long been unaware of the overlap feature and used the molecular clock, which is only good for the linear equidistance, to explain the maximum equidistance.

The molecular clock interpretation of the maximum genetic equidistance result is really about the constant rate of complexity increases. People since Aristotle have long appreciated the direction of evolution towards higher complexity. Darwin’s theory has long denied this but only by ignoring inconvenient facts including the genetic equidistance phenomenon. The evidence for complexity increase is commonplace and easy to notice by common sense. The first molecular evidence for it is the maximum genetic equidistance phenomenon. What is most striking is the nearly constant rate as measured in years of the complexity increase, which can be quantitatively indicated by the fraction of non-changeable positions in a protein or the fraction of identical residues between human and a lower complexity species (Fig. 2).

**Figure 2.**
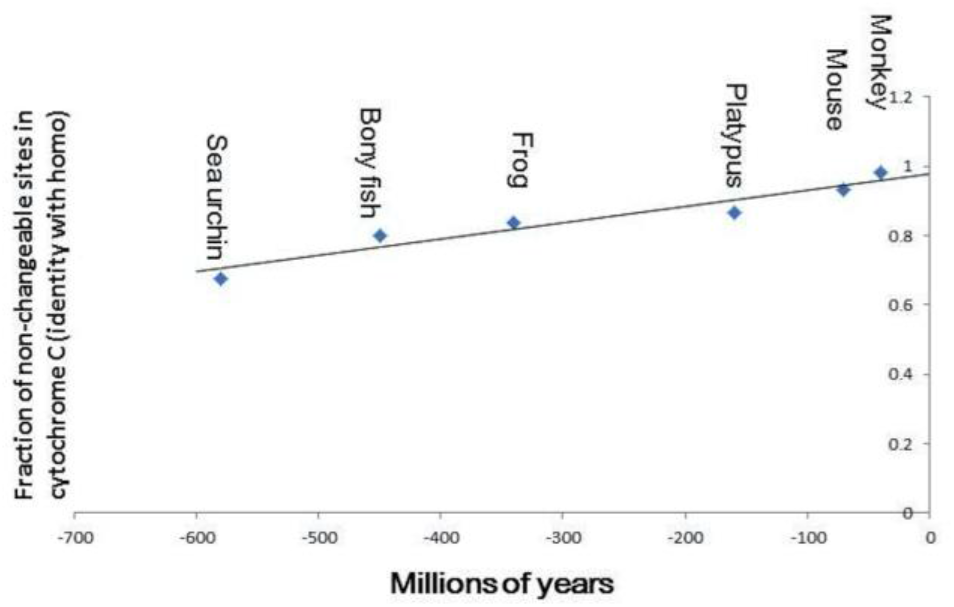
The constant rate of complexity increase. The fraction of identical residues between human and a lower complex species is equivalent to the fraction of non-changeable sites in the lower complexity species. The fraction of identical residues in cytochrome C (identity divided by length) between human and each of the species listed in the figure is plotted against the separation time between human and each of the listed species. Data for plots were obtained using homo cytochrome C to BLASTP Genbank.

As nature is written in the language of mathematics, it would be most unusual if a fundamental natural phenomenon, i.e., the constant rate of evolution towards higher complexity as measured in years, has no counterpart in mathematics and vice versa. An intriguing analogy is the pattern of prime numbers. The cumulative increase in prime numbers along the progression in natural numbers is well known to follow a nearly constant rate (Fig. 3) (Bombieri, 2000; Derbyshire, 2004; du Sautoy, 2003; Edwards, 1974; Huang, 2008; Riemann, 1859; Sabbagh, 2003). Here the progression in natural numbers is like a time clock, rigid and predictable. The appearance of prime numbers is discontinuous like a staircase and unpredictable but follows nonetheless a well defined function Li(N) as shown by the Riemann hypothesis, widely known as the most important unproved problem in mathematics (Bombieri, 2000; Derbyshire, 2004; du Sautoy, 2003; Edwards, 1974; Huang, 2008; Riemann, 1859; Sabbagh, 2003).

**Figure 3.**
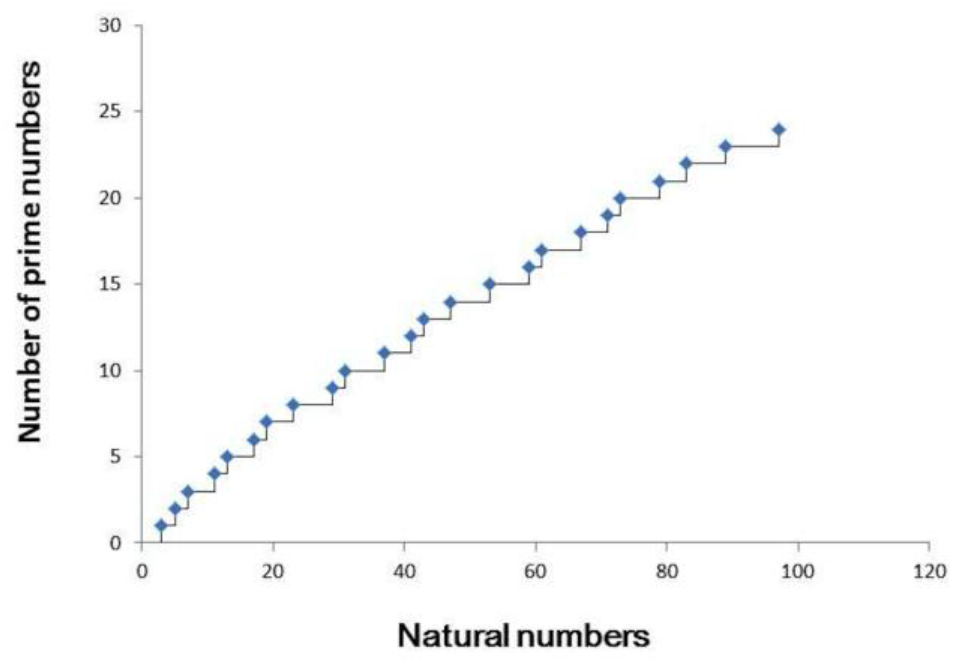
The prime number staircase. The graph counts the cumulative number of primes up to 100.

Each new appearance of a more complex species is like a new prime number, unpredictable, discontinuous, and yet constant. Individual species are well known to appear in the fossil record abruptly as evidence for the punctuated equilibrium model of macroevolution has shown (Gould and Eldredge, 1993). Defying the Darwinian gradualistic worldview, there have always been researchers since the time of Darwin who have appreciated the discontinuous nature in macro-evolutionary changes (Bateson, 1894; Denton, 1986; Denton, 2016; Forsdyke, 2011; Goldschmidt, 1940; Gould and Eldredge, 1993; Huang, 2016; Jenkin, 1867). However, the discontinuous appearance of species of higher and higher complexity still follows a very smooth and regular pattern as shown by the equidistance phenomenon. We speculate that the mystery behind the constant rate of complexity increase in nature might well turn out to be the same as that behind the constant appearance of prime numbers. Indeed, the common speculative and unproven answer to both mysteries has long been random forces.

## Acknowledgements

We thank Michael Denton for critical reading of the manuscript.

